# Inherently reduced expression of ASC restricts caspase-1 processing in hepatocytes and promotes *Plasmodium* infection

**DOI:** 10.1101/2023.07.10.548445

**Authors:** Camila Marques-da-Silva, Rodrigo P. Baptista, Samarchith P. Kurup

## Abstract

Inflammasome-mediated caspase-1 activation facilitates innate immune control of *Plasmodium* in the liver, thereby limiting the incidence and severity of clinical malaria. However, caspase-1 processing occurs incompletely in the hepatocytes and precludes the generation of mature IL-1β or IL-18, unlike in other cells. Why this is so, or how it impacts *Plasmodium* control in the liver has remained unknown. We show that an inherently reduced expression of the inflammasome adaptor molecule ASC (apoptosis-associated speck-like protein containing CARD) is responsible for the incomplete proteolytic processing of caspase-1 in hepatocytes. Transgenically enhancing ASC expression in hepatocytes enabled complete caspase-1 processing, enhanced pyroptotic cell-death, maturation of the proinflammatory cytokines IL-1β and IL-18 that was otherwise absent, and resulted in better overall control of *Plasmodium* infection in the liver mice. This however impeded the protection offered by live-attenuated anti-malarial vaccination. Tempering ASC expression in macrophages on the other hand resulted in incomplete processing of caspase-1. Our work shows how caspase-1 activation and function in host cells are fundamentally defined by ASC expression and offers a potential new pathway to create better disease and vaccination outcomes by modifying the latter.

## Introduction

Malaria, caused by *Plasmodium* parasites, remains an unresolved global health burden that impacts more than half of the world’s human population. Upon inoculation into its mammalian hosts as sporozoites, *Plasmodium* undergoes obligatory replication and development in the hepatocytes, where they transition into the merozoite stage that infects the red blood cells (RBCs). Infection of the RBCs would initiate the blood stage of malaria which is responsible for almost all of the morbidity and mortality associated with this disease. The interventions that block the progression of *Plasmodium* to its blood stage are therefore expected to prevent clinical disease and transmission (1). However, our incomplete understanding of the fundamental biology of *Plasmodium* infection in the liver has prevented us from effectively targeting the parasites during this stage of its life cycle.

It is well established that innate immune responses generated in the liver can control *Plasmodium*, potentially limiting the incidence and severity of clinical malaria (2–6). We have shown that the inflammasome pathway is a key component in this process, wherein the receptor, absent in melanoma (AIM) 2 detects *Plasmodium* DNA and induces inflammasome-mediated caspase-1 activation and pyroptotic cell death in the infected hepatocytes (2). This process is instrumental in controlling *Plasmodium* infection in the liver and limiting its progression to the blood-stage, as well as, offering Plasmodium antigens to the antigen-presentation machinery in the liver (2–4).

Inflammasomes are large macromolecular complexes generated in the host cell cytosol following the detection of pathogen-associated molecular patterns (PAMPs) such as DNA by intracellular pattern recognition receptors (PRRs). The inflammasomes are typically composed of such PRRs, the adaptor molecule, ASC (apoptosis-associated speck-like protein containing CARD (caspase recruitment domain)) when the PRRs do not possess their own CARD, and the zymogen procaspase-1. Procaspase-1 undergoes proximity-driven autoproteolysis at such inflammasome complexes to its constituent CARD, p20, and p10 subunits, following which, the p20 and p10 subunits heterodimerize to generate enzymatically active caspase-1 (7). Caspase-1 formed in this manner is believed to proteolytically activate the membrane pore-forming molecule, Gasdermin D (GSDMD), and the proinflammatory cytokines IL-1 and IL-18 (7–10).

It is noteworthy that the above paradigm was established via studies conducted primarily in the cells of myeloid lineage (7). The dynamics of procaspase-1 processing is quite distinct in the hepatocytes, wherein, they undergo only partial proteolysis into a p32 caspase-1 species composed of unseparated p20 and p10 subunits (2). Although p32 activates GSDMD and induces pyroptotic cell death following exposure to PAMPs, it does not facilitate the maturation of IL-1 or IL-18 in the hepatocytes, much like when caspase-1 p32 was artificially generated by mutating the procaspase-1 gene in the bone-marrow-derived macrophages (BMDMs) (2, 11). The molecular mechanism responsible for limiting procaspase-1 processing to the p32 form in hepatocytes has remained unknown. Filling this fundamental knowledge gap is critical in improving our current understanding of the innate immune responses in the liver.

Innate immune responses such as inflammasome-mediated caspase-1 activation are also key drivers of downstream adaptive immune responses (12). Caspase-1 function is necessary to recruit antigen-presenting cells (APCs) that prime protective CD8 T cell responses against hepatotropic infections such as *Plasmodium* colonizing the liver, following natural infections or vaccinations (4, 13–16). Determining why caspase-1 processing is limited in the hepatocytes would also pave way to identifying potential new intervention strategies to enhance adaptive immune responses against pathogens infecting the liver such as *Plasmodium*.

In this work, we show that procaspase-1 is terminally processed to the p32 caspase-1 species owing to the inherently reduced expression of ASC in the hepatocytes. Transgenic overexpression of ASC resulted in the maturation of procaspaspase-1 into p20 and p10, generation of mature IL-1β and IL-18, and enhanced GSDMD-mediated pyroptotic cell death in the hepatocytes. This also resulted in significantly better innate immune control of liver-stage malaria. In addition to advancing our basic understanding of liver biology, our discovery is expected to have key implications in anti-malarial drug and vaccine designs.

## Materials and Methods

### Mice and pathogens

C57BL/6 (B6) mice were purchased from the Jackson Laboratory. All mice were housed with appropriate biosafety containment at the animal care units of the University of Georgia and the animals were treated and handled in accordance with the guidelines established by it. *Anopheles stephensi* mosquitos infected with *P. berghei ANKA* (*Pb*) and *P.berghei* expressing luciferase (*Pb-Luc*) were reared at the SporoCore insectary facility at the University of Georgia.

### Primary mouse hepatocyte culture, *in vitro* sporozoite infection, and PRR stimulation

Primary hepatocytes were isolated from mice as described in detail before (4). In short, the inferior vena cava of anesthetized (2,2,2-Tribromoethanol, 300 mg/Kg)) mice were catheterized (BD auto guard, 22G) aseptically to perfuse the liver by draining through the hepatic portal vein. Steady-state perfusion of the liver was performed first with PBS (4ml/ min for 5 minutes), then Liver Perfusion Medium (4ml/min for 3 minutes, Gibco), and finally the Liver Digest Medium (4ml/ minute for 5 minutes, Gibco). The digested liver was excised, single cell suspension made, and resuspended in a wash solution of 10% fetal calf serum (FCS, Sigma-Aldrich) in DMEM (Gibco). Hepatocyte fraction was recovered by centrifugation at 57g, from which the debris and the dead cells were removed by density gradient centrifugation with a 46% Percoll (GE Healthcare) gradient. The remaining cells were counted and resuspended in DMEM with 10% FCS. Typically, 6×10^5^ of these cells-the hepatocytes-were plated on flat-bottomed collagen-coated plates and incubated at 37°C, 5% CO_2_. 6-8×10^5^ primary mouse or human (obtained from BioIVT) hepatocyte cultures were infected with 2-4×10^4^ sporozoites in each well of a 6-well plate. Cultures were further incubated for the desired time to allow infection and liver-stage parasite development. Hepatocytes were stimulated with LPS (100 ng, Invivogen) and ATP (5mM, Sigma) for 16 hours (h). Disulfiram was used as a pre-treatment *in vitro* in some experiments, at 100nM for 6h before LPS and ATP addition. 3×10^6^ BMDMs were treated with LPS (3.5h) followed by 5mM ATP (0.5h) as described in detail before (17). Culture supernatants and/or cell lysates were obtained at various time points post-infection (p.i.) or post-stimulation as indicated.

### Determining differential transcription

Distinct datasets were built for mice and humans. Mice RNAseq Illumina reads were retrieved from NCBI Sequence read archive (SRA) under accessions SRR9961636, SRR9961637, SRR9961638, SRR9961644, SRR9961945, and SRR9961646 for hepatocytes, and SRR6246979, SRR5714120, and SRR5714119 for macrophages. The human hepatocytes data used were from our published single-cell RNAseq results (2). Data on monocytes were retrieved from SRA under accessions SRR12539860, SRR12539861, and SRR12539862. The Illumina reads were aligned against the NCBI reference genome for mice (GRCm38) or Humans (GRCh38) using HISAT2 v2.1.0(18). The alignment files were then sorted by SAMtools v.1.6(19) and submitted to HTseq v0.9.1(20) to generate the count file. The count file was then used for the differential expression analysis under R, using DEseq2 (21). The data were normalized and filtered for p values of <0.01.

### Macrophage differentiation from bone-marrow

BMDMs were prepared as described in detail previously (17). In short, the bone marrow cells were grown in L-cell-conditioned IMDM medium (ThermoFisher) supplemented with 10% FCS, 1% non-essential amino acids, and 1% penicillin-streptomycin for five days to differentiate into macrophages. On day 5, BMDMs were seeded in 6-well cell culture plates. The next day, the BMDMs were ready for use.

### *Plasmodium* inoculations in mice

For sporozoite challenge experiments to determine liver-parasite loads, salivary glands of parasitized *A. stephensi* mosquitoes were dissected and the sporozoites were isolated, counted, and injected 2×10^4^ in 200μL RPMI with 1% mouse serum (Innov-research) retro-orbitally as described in detail before (22).

### Flow Cytometry

Cell obtained from the liver were stained with anti-CD45, F4/80, CD11c, or CSF1R (Biolegend) to determine the frequencies and counts of the myeloid cells as presented in detail before (4). To determine CD8 T cell responses, blood was collected from mice through tail or retro-orbital bleeding, the RBCs lysed with ACK lysis buffer, and the leukocyte fraction stained. The cells were stained for surface markers with appropriate antibodies or tetramers in PBS for 30 minutes prior to washing and resuspending in PBS to analyze by flow cytometry as described in detail before (4, 23, 24). CSF1R^+^ APCs were designated as CD45^+^ F4/80^+^ CD11c^+^ CSF1R^+^ cells and Plasmodium-specific CD8 T cells were designated as GAP50 tetramer^+^ CD8 T cells. Data were acquired on a Quanteon flow cytometer (Agilent) and analyzed with Flowjo (Treestar).

### Assessment of liver parasite burden

Liver parasite burden was assessed by quantitative real-time RT-PCR for parasite 18s rRNA in the livers of mice challenged with sporozoites isolated from infected mosquitoes as described before (25). Total RNA was extracted from the liver at the indicated time points after *Plasmodium* infection using TRIzol (Sigma), followed by DNase digestion and clean up with RNA Clean and Concentrator kit (Zymo Research). 2μg liver RNA per sample was used for qRT-PCR analysis for *Plasmodium* 18S rRNA using TaqMan Fast Virus 1-Step Master Mix (Applied Biosystems). Data were normalized for input to the GAPDH control for each sample and are presented as ratios of *Plasmodium* 18s rRNA to GAPDH RNA signals. The ratios depict relative parasite loads within an experiment and do not represent absolute values. *Pb-Luc* was used to assess the kinetics of replication and clearance of *Plasmodium* infection in the liver of mice. For bioluminescent detection, mice were injected with D-luciferin (150 mg/kg; SydLabs) intraperitoneally 5 minutes prior to being anesthetized using 2% (vol/vol) gaseous isoflurane in oxygen and imaged using an IVIS 100 imager (Xenogen). The quantification of bioluminescence and data analysis was performed using Living Image v4.3 software (Xenogen).

### Cell-death assay

Propidium Iodide (PI) uptake assay for pyroptosis: PI staining distinguishes programmed cell-death from whole cell-lysis (26). To determine the extent of pyroptotic cell death, hepatocytes transfected with designated plasmids were seeded at 2×10^4^ cells per well of an opaque-wall, clear-bottom 96-well plate (Corning) overnight. Culture media was replaced with DMEM without phenol red and cells received the specified treatments or infections. All culture wells were supplemented with 6 mg/ml PI (Invitrogen) and were incubated at 37°C for 40 minutes. Subsequently, the cells were washed thrice with PBS and, the plate was sealed with a clear, adhesive optical plate seal (Applied Biosystems). The frequencies of PI-stained cells were estimated using a plate reader (Molecular Devices SpectraMax iD3) and 100% cytolysis was calculated. The maximum signal was determined following treatment with 1% triton-x. Data were presented as experimental group PI signal / maximum PI signal in percentage, and normalized using untreated sample levels. Untreated hepatocyte cultures over-expressing ASC or eGFP exhibited no discernable differences in basal PI staining levels.

### Transfection of cells

Hepatocytes (7×10^5^ cells) were transfected with various plasmids (at 6 μM or indicated concentrations) using the Mouse/ Rat hepatocyte Nucleofector kit (Lonza) following the manufacturer’s protocol, yielding approximately 50% transfection efficiency in the primary cells. For overexpression of ASC, pcDNA3-N-Flag-ASC1 (Addgene, #75134) plasmid was used. For overexpression of procaspase-1, pcDNA3-N-Flag-Caspase-1 (item #75128) was used. The transfected cells were transferred to collagen-coated wells 16h prior to infection or the treatments. For silencing, cells were transfected with ON-TARGETplus SMARTpool siRNAs (Dharmacon). Differentiated BMDMs were detached by treatment with trypsin and resuspended at a concentration of 10^6^ cells per 100 μl or nucleofector solution. These cells were transfected with ON-TARGETplus SMARTpool siRNAs (Dharmacon) using the Mouse Macrophage Nucleofector kit (Amaxa) following the manufacturer’s protocol. The transfected cells were transferred to plates and allowed to recover for 24-48h prior to infections or treatments.

### Microscopy

For imaging, hepatocytes were cultured overnight in collagen-coated chamber slides (ibidi), transfected with ASC-encoding plasmid, treated with LPS+ATP as described above, and stained for FLICA as per the manufacture’s protocol using the FAM-FLICA-Caspase-1 (YVAD) assay kit (Immunochemistry), and visualized using fluorescence microscopy (Keyence). In short, the adherent cells were incubated for 1h at 37°C, washed twice with 1x apoptosis buffer and stained with Hoechst stain. The fluorescence signal intensity of the FLICA-stained inflammasome complexes, as well as, the area was determined using ImageJ software from the acquired images and presented as mean signal intensity, calculated as background corrected mean fluorescence /area (27).

### Therapeutic regimen

The following treatment regimen was used in this study: Disulfiram (Sigma-Aldrich): 50mg/kg in sesame oil, i.p

### Adeno-associated viral vectors

Adeno-associated viral vectors were generated by VectorBuilder Inc. by encoding the genes of interest under the control of the albumin promoter to limit target protein expression to the hepatocytes (28). AAV-DJ strain was used as the background for efficient transduction of hepatocytes *in vivo* (29). Virus stocks were resuspended in PBS and inoculated intravenously at 1×10^11^ GC/ mouse. Experiments were performed 10-14 days after inoculation for efficient transduction and protein expression.

### ELISA

To quantify specific protein levels in culture supernatants of *ex vivo* cultured hepatocytes, ELISA was performed as described in detail before (30). In short, serial dilutions of supernatants or whole cell lysates (normalized for total protein) were coated in triplicate in 96-well format in 0.1M carbonate bicarbonate buffer (pH 9.6) overnight at 4°C, washed thrice with 0.05%Tween20 in PBS

(PBS-T), blocked for 60 min with 1%BSA/PBS, probed with anti-mouse anti-caspase-1 (Clone Casper-1, Adipogen), anti-ASC (F-9, Santa Cruz), anti-IL-1β (polyclonal, R&D) or anti-IL-18 (Clone 125-25, MBL international) as applicable for 60 min at 37°C, washed 5 times with PBS-T, probed with the corresponding HRP conjugated secondary antibodies in 1% BSA/PBS and again washed 3 times with PBS-T before developing. All ELISA assays were developed using TMB liquid substrate system (Sigma), stopped with 2N sulfuric acid, and then read at 450nm using an ELISA microplate reader (Molecular Devices SpectraMax iD3).

### Western blot

Western blots were performed as described before (2, 17). In short, the cells in culture were lysed along with the supernatant using RIPA Buffer or sample loading buffer containing dithiothreitol (DTT) and sodium dodecyl sulfate (SDS) at the indicated time points. The supernatant was included in the assays inducing Caspase-1 activation since caspase-1 activation and cell-death would release the cytosolic contents into the supernatant. The protein samples were run on 12% SDS-polyacrylamide gel by electrophoresis and transferred to PVDF (Millipore) or Nitrocellulose (BioRad) membranes. After blocking with 5% skimmed milk or Odyssey blocking Buffer (Licor) for 1h at room temperature, the membranes were probed with the primary antibodies: mCaspase-1p20 (Clone Casper-1, Adipogen), mCaspase-1p10 (Clone Casper-2, Adipogen), mIL-1β (Clone D3H1Z, Cell Signaling), mIL-18 (Clone 39-3F, MBL), mGasdermin D (Clone EPR19828, AbCam), or mASC (F-9, Santa Cruz) at 4^0^C overnight, washed with tris-buffered saline containing 0.1% Tween-20 (TBST) four times and incubated at room temperature for 45 min with polyclonal secondary anti-rabbit, anti-rat or anti-mouse chemiluminescent (Jackson Immunoresearch) or IRDye-conjugated (LICOR) antibodies as applicable. After 4 washes with TBST, the proteins were visualized using a chemiluminescence detection reagent (Millipore) or directly by fluorescence (LICOR). Loading controls (LC) reflect the total amount of protein in the specified lane and rely on the housekeeping protein, beta-tubulin when whole-cell lysates were analyzed. When culture supernatants also constituted the samples, an arbitrary protein band in the corresponding SDS PAGE acted as the LC, as published before (2, 31, 32) and is known to provide a more precise comparison across samples (33). Relative band densities were not averaged across replicate blots since the signal intensities representing same amount of protein vary drastically with varying imaging conditions such as the exposure time, the detection systems used etc., and is known to be non-linear when infrared or chemiluminescence detection methods are applied (34, 35).

Chemical striping of the western blot membranes when employed followed the manufacturer’s protocol (LICOR). In short, membranes were incubated for 7 minutes in fluorescent striping buffer, washed 3 times in PBS and re-probed with the primary antibodies. Band density quantification reflects the relative protein amounts as a ratio of each protein band signal intensity to the lane’s loading control, determined using Image J software.

### Statistical analyses

Data were analyzed using Prism7 software (GraphPad) and as indicated in the figure legends.

## Results

### Incomplete processing of caspase-1 in hepatocytes

Based on studies in BMDMs, caspase-1 activation is believed to occur through autoproteolysis of procasaspe-1 into its constituent CARD, p20, and p10 subunits, following which, the p20 and p10 subunits heterodimerize to generate enzymatically functional caspase-1 (**Fig S1A**). In contrast, as we have shown previously in both human and mouse hepatocytes, procaspase-1 is terminally processed to a p32 caspase-1 species composed of unseparated p20 and p10 subunits (**Fig S1A**) (2). Caspase-1 p32 species has been consistently observed in primary murine hepatocyte cultures infected with *P. berghei* (*Pb*) or treated with the standard inducers of the inflammasome pathway, LPS+ATP, and for up to 24 hours (h) post-exposure (**Figure S1B-C**), indicating that it is a stable protein species (2). Why caspase-1 processing is naturally limited to its intermediate p32 form in the hepatocytes remained unknown.

### Inherently reduced baseline expression of ASC in hepatocytes

We considered the possibility that the incomplete processing of caspase-1 in hepatocytes is due to alternate splicing of procaspase-1, making the interdomain linker (IDL) connecting the p20 and p10 domains not amenable to proteolytic cleavage. Nevertheless, bioinformatic analysis of our (4) or others’ (36–39) published transcriptomic data did not reveal any procaspase-1 splice variants in human or mouse hepatocytes. Furthermore, no detectable alternately spliced transcripts of procaspase-1 were detected in either naive, *Plasmodium-*infected, or LPS+ATP-stimulated primary murine hepatocytes (2).

The adaptor molecule ASC is critical for seeding the inflammasome platform (also called ASC specks) that assimilates procaspase-1 molecules locally in the host cell cytoplasm, imparting it a catalytically active quaternary structure and autoproteolytic activity (7, 27, 40, 41). Examination of our (4) and other publicly available transcriptomic data (36–39) indicated that both procaspase-1 and ASC transcripts are inherently less abundant in both mouse and human hepatocytes when compared to the cells of myeloid lineage in either species (**Figure 1A**). This was corroborated by their baseline protein expression as well (**Figure 1B, C**). A lower overall expression of procaspase-1 would result in fewer procaspase-1 molecules being recruited to the inflammasome complex, impeding its processing. A low expression of ASC would, on the other hand, result in its sub-optimal oligomerization, leading to lower assimilation of procaspase-1 at the inflammasome complexes (42, 43). Therefore, the reduced expression of procaspase-1 or ASC were potential explanations for the suboptimal processing of proaspase-1 in hepatocytes.

**FIGURE 1:**
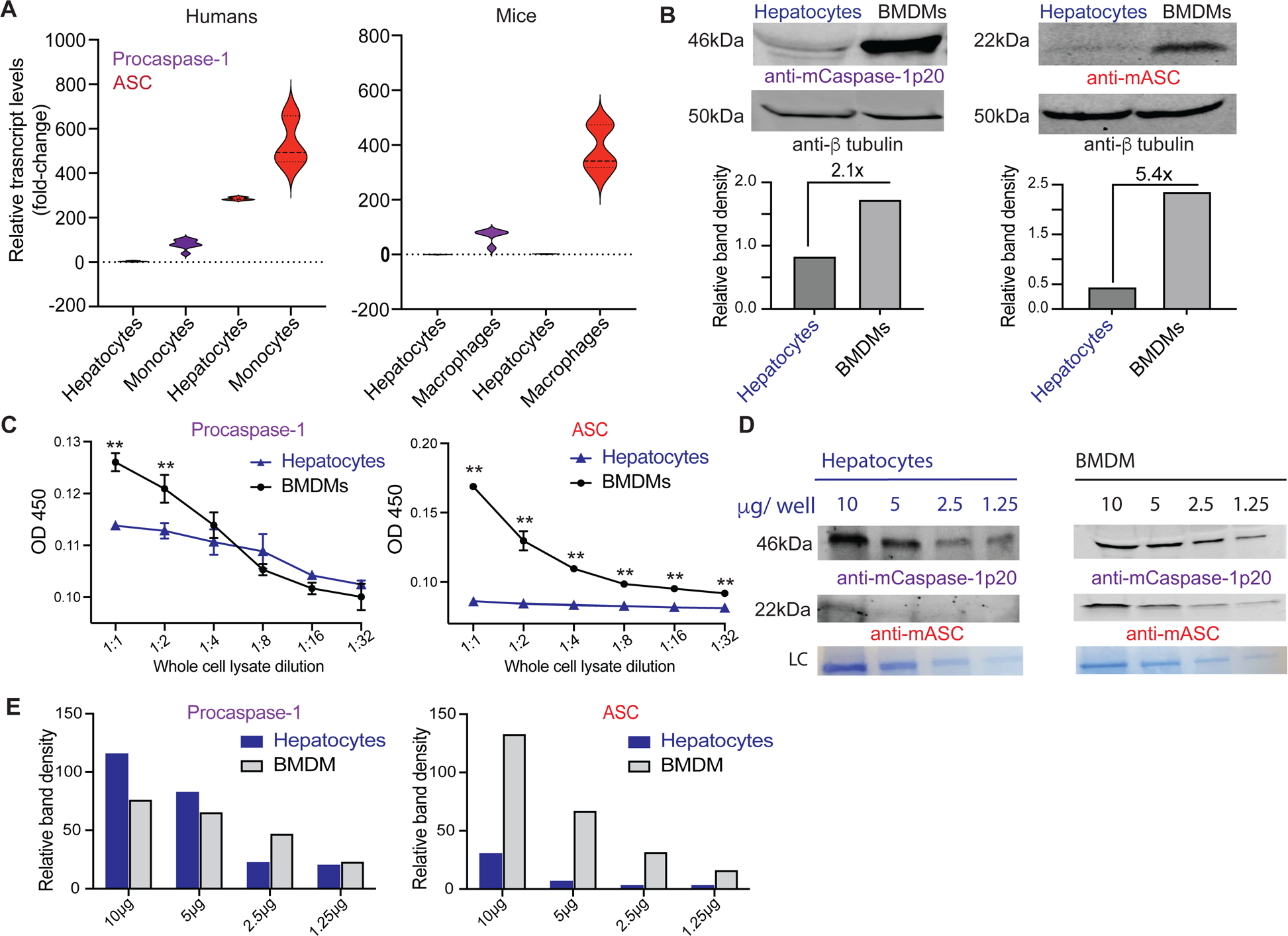
Inherently reduced expression of procaspase-1 and ASC in hepatocytes. (**A**) Combined data depicting the relative transcript levels of procaspase-1 and ASC in the human or murine primary hepatocytes or myeloid cells. (**B**) Immunoblot analysis depicting the relative expression of procaspase-1 and ASC in resting primary mouse hepatocytes or BMDMs. Bar graphs in the lower panel indicate the relative band densities of procaspase-1 or ASC normalized to beta-tubulin used as loading controls (LC) in the indicated blots. Data shown from 1 of 3 separate replicate experiments. (**C**) Relative levels of procaspase-1 (left) and ASC (right) determined by ELISA in whole-cell lysates of resting murine hepatocytes or BMDMs. Data presented as mean ± s.e.m analyzed with t-tests at each dilution of total cell lysates from three biological replicates. Data shown from 1 of 3 separate replicate experiments. *p≤0.05, **p≤0.01. (**D**) Immunoblot analysis for procaspase-1 or ASC in serially diluted whole-cell lysates of primary mouse hepatocytes or BMDMs treated with LPS+ATP for 16h or 4h (LPS 3.5h + ATP 0.5h) respectively. The blots were first probed with caspase-1-specific antibodies followed by chemical stripping and probing with ASC-specific antibodies. LC: loading control. (**E**) Quantification of the band densities of the blots shown in (**D**) depicting the relative expression of procaspase-1 (left) or ASC (right) in hepatocytes and BMDMs. (**D-E**) Representative data from 1 of 4 replicate experiments shown.

Stimulation with PAMPs such as LPS is known to increase the expression of intracellular PRRs and procaspase-1 in carious cell types (44, 45). Therefore, to determine if the reduced expression of procaspase-1 and ASC remained intact in the context of PAMP-stimulation in the hepatocytes, we treated both hepatocytes and BMDM controls with LPS+ATP. The difference in procaspase-1 expression between hepatocytes and BMDMs became less pronounced following PAMP stimulation. But, the expression of ASC continued to remain reduced in the hepatocytes despite LPS stimulation (**Figure 1D, E**). Together, these findings indicated that the baseline expression of ASC in hepatocytes is naturally lower in comparison to in the myeloid cells.

### The extent of caspase-1 processing in hepatocytes is limited by ASC expression

The formation of dense ASC specks that bring procaspase-1 molecules in close proximity to each other in the cytoplasm is a critical determinant of efficient caspase-1 processing and function, (27, 42). We predicted that transgenically enhancing the expression of procaspase-1 and/ or ASC in hepatocytes would facilitate the formation of compact inflammasome complexes, leading to complete procaspase-1 processing, resulting in the generation of caspase-1 p20 subunit. To test this hypothesis, we over-expressed procaspase-1 or ASC in primary murine hepatocytes (**Fig 2A, B**). Over-expression of ASC, but not procaspase-1 in primary murine hepatocytes generated discernable caspase-1 p20 cleavage product following the stimulation with LPS+ATP (**Fig 2C**). Enhanced green fluorescent protein (eGFP) served as the control for transgenesis. Over-expression of ASC also generated caspase-1 p20 following *Pb* infection (**Fig 2D**). Notably, the overexpression of ASC also facilitated the generation of higher p32 levels in such hepatocytes, possibly due to the higher overall inflammasome mediated caspase-1 activation in such cells.

**FIGURE 2:**
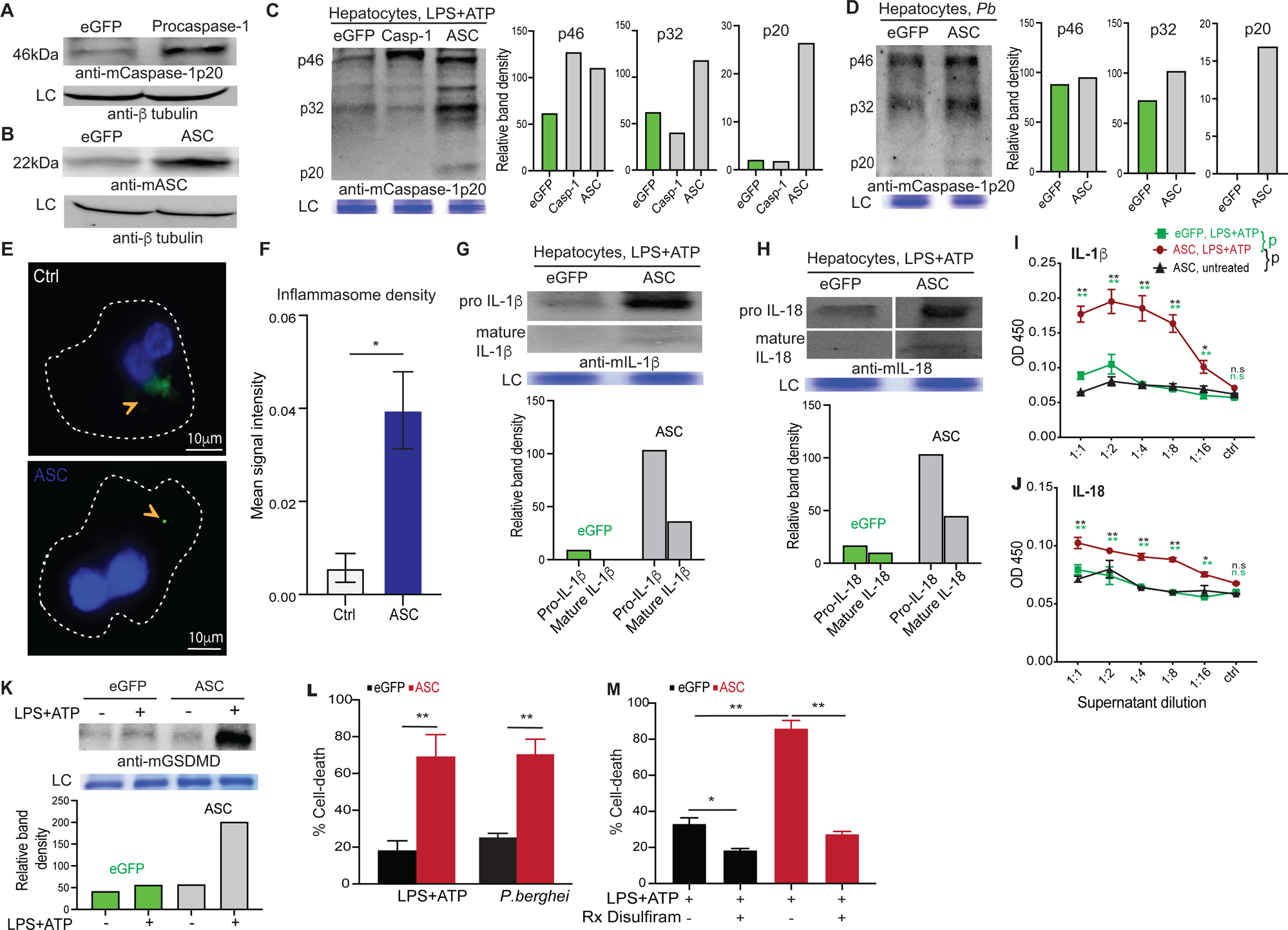
Augmenting ASC expression in hepatocytes induce complete caspase-1 processing, maturation of IL-1β and IL-18, and enhanced cell death. (**A**) Immunoblot analysis depicting the relative expression of procaspase-1 in primary mouse hepatocytes transfected with control eGFP or murine procaspase-1 genes, 24h post transfection. Representative blot from 3 separate experiments shown. (**B**) Immunoblot analysis depicting the relative expression of ASC in primary mouse hepatocytes transfected with the eGFP or murine ASC genes, 24h post transfection. Representative blot from 3 separate experiments shown. (**A-B**) Total β tubulin in cell-lysates served as loading controls. (**C**) Immunoblot analysis of caspase-1 processing in primary mouse hepatocytes transfected with eGFP, murine procaspase-1 (Casp-1), or murine ASC genes and treated with LPS+ATP for 16h. The cultures were treated starting at 24h post transfection. Representative blot from 3 separate experiments shown. Bar graphs on the right indicate relative band densities normalized to the loading controls. (**D**) Immunoblot analysis of caspase-1 processing in primary mouse hepatocytes transfected with eGFP or murine ASC genes and infected with *Pb* sporozoites 24h post transfection and examined at 16h p.i. Representative blot from 2 separate experiments shown. The bar graphs on the right indicate relative band densities normalized to loading controls. (**E**) Representative (of >10 fields, 3 replicates) pseudo-colored confocal image depicting active inflammasome complexes (arrow) generated in LPS+ATP-treated (16h) wild-type (ctrl, upper panel) or ASC-transgenic (ASC, lower panel) *ex-vivo* cultured primary murine hepatocytes, determined using caspase-1-FLICA staining. Dotted line represents the outline of the host cell. (**F**) Bar graphs indicating the intensity of caspase-1-FLICA staining in the inflammasome complexes generated in (**E**), represented as mean signal intensity. Combined data presented as mean ± SD from ≥10 fields from 3 replicate experiments. (**G-H**) Immunoblot analysis showing the relative levels of IL-1β (**G**) or IL-18 (**H**) in the supernatants of eGFP or ASC-transgenic primary murine hepatocyte cultures treated with LPS+ATP for 16h, starting at 24h post-transfection. Bar graphs below indicate relative band densities normalized to loading controls. (**I-J**) IL-1β (**I**) or IL-18 (**J**) in the supernatants of eGFP or ASC-transgenic primary murine hepatocyte cultures treated with LPS+ATP for 16h, starting at 24h post-transfection, determined by ELISA. ELISA data presented as mean ± s.e.m at each dilution of the culture supernatant, analyzed using 2-way ANOVA with Dunnett’s correction, comparing the color-coded groups. Representative data shown from 1 of ≥2 separate experiments. (**K**) Immunoblot analysis for activated GSDMD in eGFP or ASC-transgenic primary murine hepatocytes in culture co-incubated for 16h with or without LPS+ATP, starting from 24h post-transfection. LC: loading control. Bar graphs below indicate relative band densities normalized to loading controls. Representative data shown from 1 of 3 separate experiments. (**L**) Frequency of cell-death determined by propidium iodide (PI) staining in eGFP or ASC-transgenic primary murine hepatocytes in culture treated with LPS+ATP or infected with *Pb* (16h). (**M**) Frequency of cell-death determined by PI staining in eGFP or ASC-transgenic primary murine hepatocytes in culture treated with LPS+ATP (16h) and with or without disulfiram (6h prior to stimulation). (**L,M**) Data normalized to untreated transfected cells presented as mean ± s.e.m and analyzed using ANOVA with Dunnett’s correction to yield the indicated p values. Representative data shown from 1 of 3 separate experiments. n.s: p>0.05, *p≤0.05, **p≤0.01

Enhancing ASC expression also resulted in the generation of more condensed and catalytically active inflammasome complexes in the hepatocytes, as evidenced by ASC specks with relatively higher-intensity of FLICA staining (**Figure 2E, F**). It is known that the intensity of FLICA staining directly- and the size of ASC specks inversely-correlate with inflammasome and caspase-1 activity in cells (46). Taken together, the above findings suggested that the baseline expression of ASC in host cells may be a determinant of the extent of caspase-1 activation. This meant that tempering ASC expression in BMDMs would result in incomplete processing of caspase-1 and the generation of p32 in BMDMs following PAMP stimulation. To test this, we transfected BMDMs with increasing doses of siRNA to tune down ASC expression without completely eliminating it (**Figure S2A**). While fully abrogating the expression of ASC prevented any caspase-1 activation in BMDMs following LPS+ATP stimulation (**Fig S2B**), tempering it resulted in the progressive appearance of p32, with concurrently diminishing p20 levels (**Fig S2C, D**). These observations further confirmed that the level of ASC expression in host cells is a key determinant of the extent of procaspase-1 processing in the context of ASC-mediated inflammasome activation.

Over-expression of ASC in primary murine hepatocytes also generated detectable levels of mature IL-1β and IL-18 (**Figure 2G-H**), as well as, higher levels of active GSDMD in response to LPS+ATP stimulation (**Figure 2K**). This phenotype was also accompanied by a significantly higher cell-death response in ASC-transgenic hepatocytes following LPS+ATP stimulation or *Plasmodium* infection, compared to the control eGFP over-expressing hepatocytes (**Figure 2L**). The frequency of eGFP-transgenic hepatocytes undergoing cell-death following *Pb* infection is in the range of what has been observed in wild-type hepatocytes in previous studies (2). Therapeutically blocking GSDMD function using disulfiram negated this phenotype when examined in the context of the standard LPS+ATP stimulation (**Figure 2M**). These data indicated that the enhanced pyroptosis observed in ASC-transgenic hepatocytes was potentially driven by the enhanced GSDMD activation.

The data in **Figures 2C** and **S2B** indicated that higher ASC expression was also associated with slightly higher procaspase-1 levels in the hepatocytes and BMDMs. Enhancing ASC expression elevated the levels of pro IL- and IL-18 as well (**Figures 2G, 2H**). Although procaspase-1 is believed to be constitutively expressed in cells, pro-inflammatory cytokines such as IL-1 produced as a result of complete processing of caspase-1 can enhance procaspase-1 expression through autocrine or paracrine nuclear factor kappa B (NF-κB)-mediated signaling (45, 47). Similarly, activated IL-1 and IL-18 can further induce the expression of pro IL-1 and IL-18 in a paracrine or autocrine fashion in the host cells (48). Nevertheless, the fact that reducing ASC expression in BMDMs resulted in partial processing of procaspase-1, as well as the absence of caspase-1 p20 following overexpression of procaspase-1 in hepatocytes pointed towards a direct role of ASC in regulating the extent of caspase-1 processing, plausibly independently of the altered procaspase-1 levels.

Taken together, the above data indicated that caspase-1 processing occurs incompletely in hepatocytes possibly due to the lower expression of ASC, and that enhancing ASC induced complete processing of caspase-1, production of mature IL-1β and IL-18, and improved GSDMD-mediated pyroptotic cell-death when the inflammasome pathway was induced.

### Augmenting ASC expression in hepatocytes improves protection from malaria

GSDMD-mediated pyroptotic cell death induced by the AIM2-caspase-1 axis is a key component of natural immunity to malaria in the liver (2, 3). Our ability to enhance pyroptosis in *Plasmodiu*m-infected hepatocytes by augmenting ASC expression suggested that the latter would result in better control *Plasmodium* in the liver. To test this, we generated transgenic adeno-associated virus (AAV) on the AAV-DJ background where ASC (AAV-ASC) or control eGFP (AAV-eGFP) were placed under an albumin promoter. AAV-DJ variant was employed to ensure high transduction efficiency and protein expression in hepatocytes *in vivo*, and the albumin promoter was used to restrict the expression of the target proteins to only the hepatocytes (29, 49–51). AAV-eGFP or AAV-ASC inoculated mice were challenged with luciferase-expressing *Pb* (*Pb-Luc*), and the kinetics of *Plasmodium* infection was determined (**Figure 3A**). ASC over-expression was confirmed by western blot in primary hepatocytes isolated from the AAV-ASC inoculated mice (**Figure 3B**). Transgenically enhancing ASC expression in the hepatocytes resulted in the rapid control of *Plasmodium* in mice (**Figure 3C, D**). This phenotype was however undone by disulfiram treatment (**Figure 3E**). These data suggested that increasing ASC expression in hepatocytes would result in improved innate immune control of *Plasmodium* in the liver, possibly though enhanced pyroptotic elimination of the infected hepatocytes. Of note, proinflammatory cytokines such as IL-1 are known to not have a direct role in the innate immune control of liver-stage malaria (2).

**FIGURE 3:**
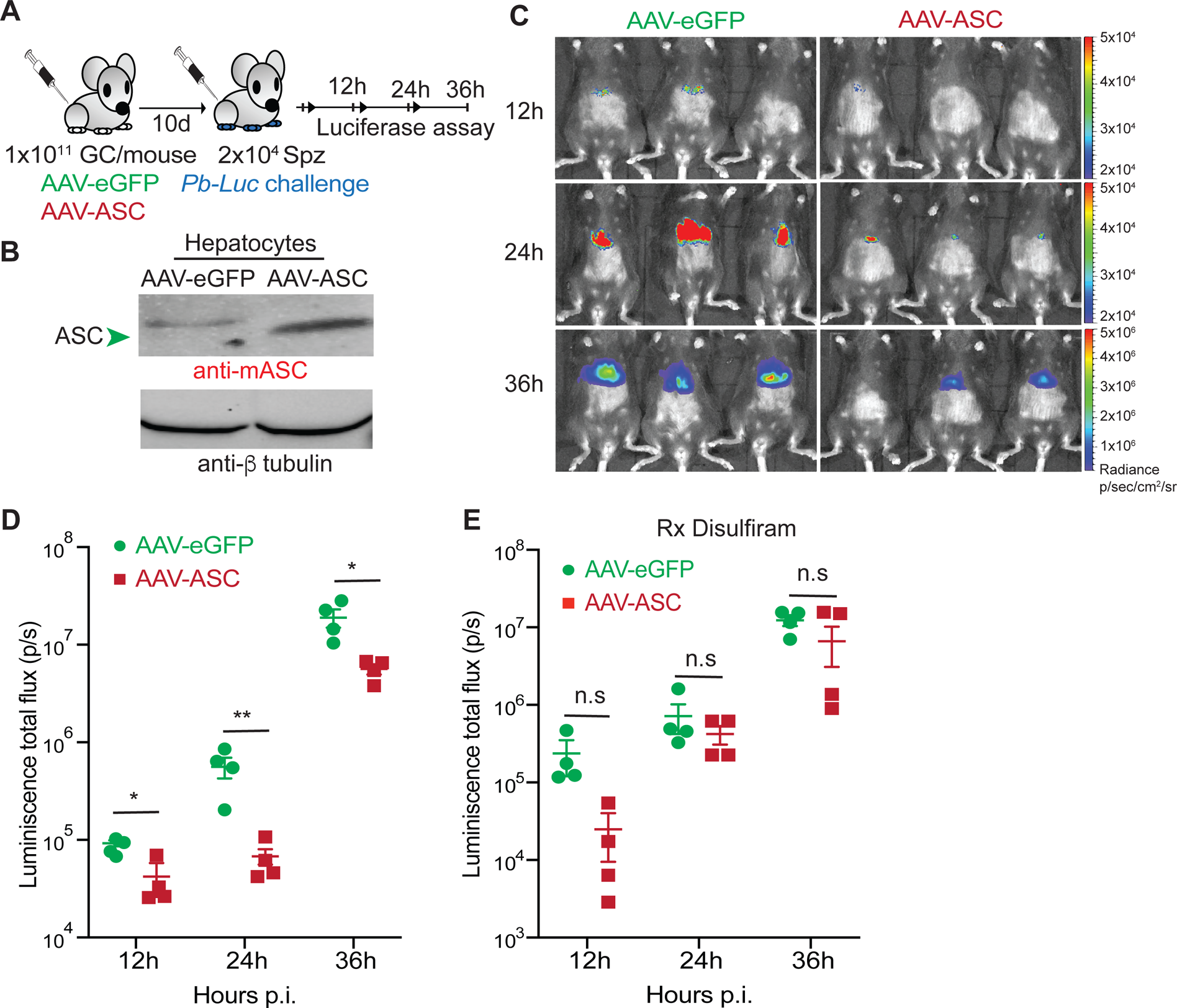
Enhancing ASC expression in hepatocytes facilitates immunity to liver-stage malaria. (**A**) Experimental scheme: 1×10^11^ genome copy (GC) of control AAV-eGFP or AAV-ASC were inoculated into B6 mice intravenously (i.v) and challenged with 2×10^4^ sporozoites of luciferase-transgenic *P. berghei* (*Pb-Luc)* i.v. at 10d post viral delivery. Mice were imaged for luminescence signal at the indicated time-points to determine the kinetics of parasite control in the liver. (**B**) Immunoblot analysis for the relative expressions of ASC in the primary hepatocytes isolated from mice inoculated with 1×10^11^ genome copy (GC) of control AAV-eGFP or AAV-ASC i.v, at 10d post viral inoculation, as depicted in (**A**). Total β tubulin in hepatocyte lysates served as loading controls. Data shown from 1 of the 2 replicate experiments. (**C**) Representative rainbow images of luminescence signal indicating liver-parasite burdens at the indicated time points in mice inoculated with AAV-eGFP or AAV-ASC, and challenged with *Pb-Luc* at 10d post viral delivery as depicted in (**A**). Representative image overlays and signal intensity scales from a total of 3 separate experiments shown. The scales are different across the time-points. (**D**) Scatter plots showing the relative parasite burdens at the indicated time points in mice inoculated with AAV-eGFP or AAV-ASC, and challenged with *Pb-Luc* at 10d post viral delivery as depicted in (**A**), with 4 mice/group. (**E**) Scatter plots showing relative parasite burdens at the indicated time points in mice inoculated with AAV-eGFP or AAV-ASC, and challenged with *Pb-Luc* at 10d post viral delivery as in Figure (**A**), with 4 mice/group. These mice were treated with disulfiram at 0 and 1d post *Pb-luc* challenge. (**D-E**) Each dot represents an individual mouse and data presented as mean ± s.e.m, analyzed using 2-tailed t-tests for each time-point, yielding the indicated p values. Representative data from 1 of 3 separate experiments shown. n.s: p>0.05, *p≤0.05, **p≤0.01.

Intriguingly, enhancing ASC expression in hepatocytes also resulted in an increased influx of total CD11c^+^ APCs, as well as, the specialized CSF1R^+^ CD11c^+^ APCs to the liver following *Plasmodium* infection (**Figure 4A-C**). This may be owing to the the increased pyroptotic cell-death and the production of IL-1 and IL-18 associated with ASC over-expression in hepatocytes (52, 53). Although not pertinent to the direct control of an ongoing infection, the monocyte-derived CSF1R^+^ CD11c^+^ APC subset is responsible for acquiring *Plasmodium* antigens from the infected hepatocytes undergoing cell-death and priming protective CD8 T cell responses (4). Such CD8 T cells are the primary mediators of protection induced by live-attenuated vaccines targeting such as the radiation or genetically attenuated sporozoite-based malaria vaccines (13, 54, 55). These findings, therefore predicted that enhancing ASC expression in hepatocytes would lead to better vaccine-induced immunity to malaria.

**FIGURE 4.**
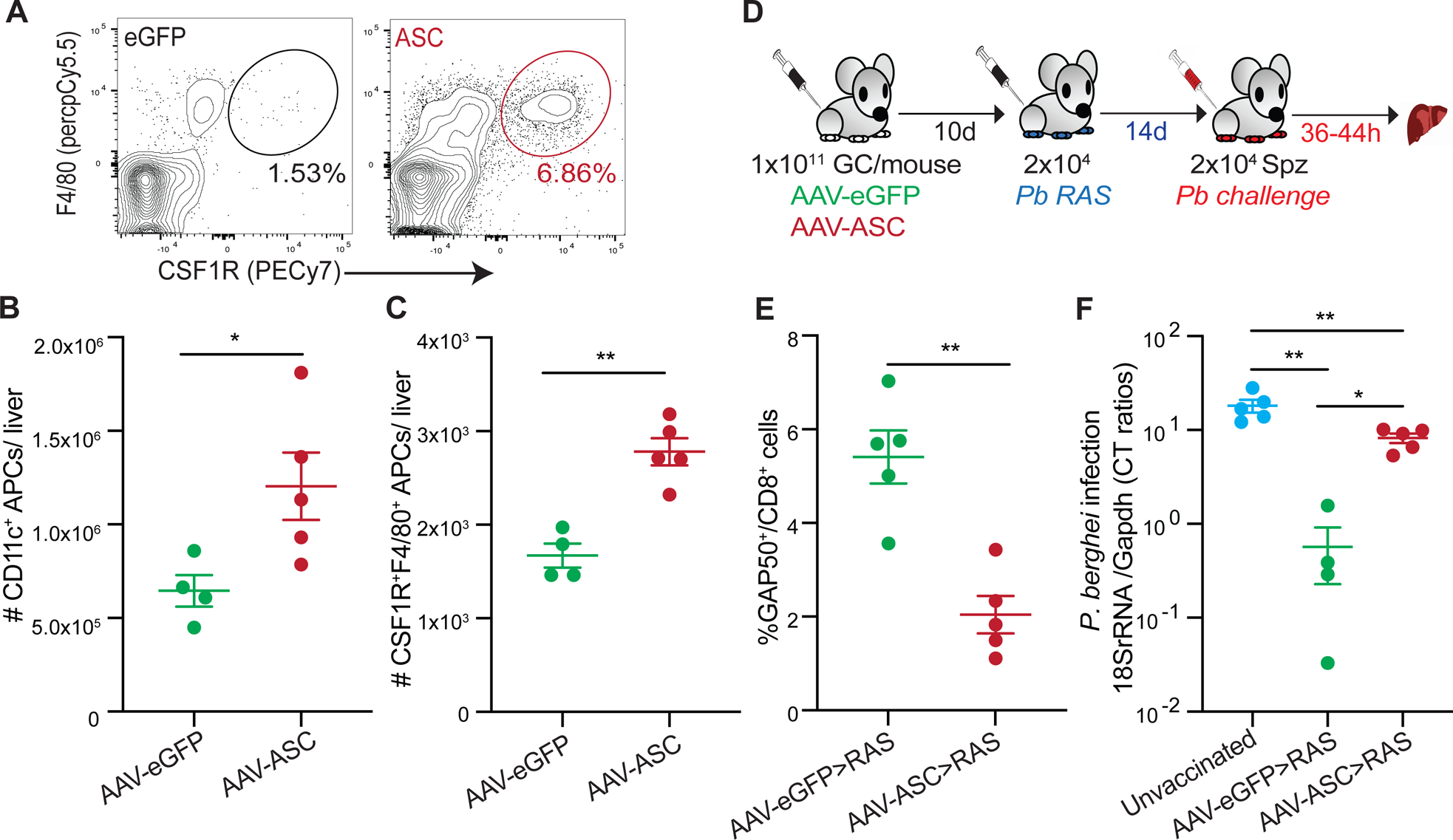
Enhancing ASC expression in hepatocytes impedes vaccine-induced immunity to malaria. (**A**) Representative flow plots depicting the frequencies of CSF1R^+^APCs infiltrating the livers of AAV-eGFP or AAV-ASC inoculated mice challenged with *Pb-Luc* at 10d post viral delivery, determined at 36h p.i. Data shown from 1 of 3 separate experiments. (**B-C**) Scatter plots depicting the total numbers of CD11c^+^ (**B**) and CSF1R^+^ CD11c^+^ (**C**) APCs in the livers of AAV-eGFP or AAV-ASC inoculated mice challenged with *Pb-Luc* at 10d post viral delivery, determined at 36h p.i. (**D**) Experimental scheme: B6 mice were inoculated with 1×10^11^ GC of control AAV-eGFP or AAV-ASC i.v. and immunized with 2×10^4^ *P. berghei* RAS (*Pb RAS*) i.v. at d10 post viral inoculation. The vaccinated mice were challenged with 2×10^4^ *Pb* sporozoites at 14d post vaccination. Blood was collected to examine CD8 T cell responses and liver collected to determine parasite loads. (**E**) Scatter plots depicting the frequencies of GAP50-specific CD8 T cells in circulation in the AAV-eGFP or AAV-ASC inoculated *Pb RAS* vaccinated mice as in (**D**) at 7d post-vaccination with *Pb RAS*. (**F**) Scatter plots depicting the relative liver parasite burdens at 44h post *Pb* challenge in AAV-eGFP or AAV-ASC inoculated mice, that were *Pb* RAS vaccinated, as depicted in (**D**). Data presented as mean ± s.e.m and analyzed using 2-tailed t-tests (**B,C,E**), or ANOVA with Tukey’s correction (**F**) to yield the indicated p values with representative data shown from 1 of 3 separate experiments with at least 3 mice/group. *p≤0.05, **p≤0.01.

To test this, we immunized mice inoculated with AAV-eGFP or AAV-ASC with radiation-attenuated sporozoites (RAS) before challenging them with virulent *Pb* sporozoites (**Fig 4D**). Surprisingly however, AAV-ASC inoculated mice induced significantly lower *Plasmodium* GAP50 epitope-specific CD8 T cell responses (**Fig 4E**). The GAP50^+^ CD8 T cell responses act as a reliable surrogate for the overall protective CD8 T cell responses generated against liver-stage malaria and is a key predictor of RAS vaccine-induced immunity in the mouse model (23). In support of the above observation, AAV-ASC inoculated mice exhibited sub-optimal protection from *Pb* challenge following RAS immunization. Although seemingly counter-intuitive, these results are in agreement with published findings that stronger innate immune responses in the liver impede sporozoite-based vaccine-induced immunity to malaria (56), and is possibly owing to the rapid elimination of RAS from the livers of AAV-ASC inoculated mice, precluding the generation of adequate protective epitopes following RAS vaccination.

These data show that enhancing ASC expression in hepatocytes induced significantly better control of *Plasmodium* infection in the liver, potentially by driving a more efficient GSDMD-mediated elimination of the infected hepatocytes. Although this enhanced the recruitment of CSF1R^+^APCs to the liver, it impeded the generation of protective CD8 T cell responses and vaccine-induced immunity to malaria.

## Discussion

The existing model for the caspase-1 activation dynamics has been derived from studies conducted primarily in cells of myeloid lineage (2, 7, 11, 27). However, caspase-1 processing that occurs in hepatocytes following infections with *Plasmodium* or stimulation with the PAMPs such as LPS+ATP used to establish the model in myeloid cells do not fit this model (2). Despite its potential implications for basic research and therapeutics pertinent to the liver, the biological reasons behind such a deviation have remained unknown. Here we show that caspase-1 processing occurs unconventionally and incompletely in hepatocytes due to the relatively lower expression of ASC. By transgenically enhancing ASC expression in hepatocytes, we enabled complete proteolysis of procaspase-1, maturation of IL-1β and IL-18, and enhanced GSDMD-mediated pyroptotic cell-death. Reducing ASC expression in myeloid cells on the other hand resulted in incomplete processing of procaspase-1. These findings suggested that ASC expression is a key regulator of the extent of caspase-1 processing in host cells. While increasing ASC in hepatocytes engendered better innate immune control of liver-stage malaria, it impeded the generation of *Plasmodium*-specific CD8 T cell responses and vaccine-induced protection against malaria.

The liver is an important barrier organ that mounts robust immune responses against a variety of pathogens, toxins, and allergens which access our body (57–59). The ability of the liver to maintain an immunotolerant environment is however critical for its functional integrity (57–61). Although home to a large collection of specialized immune cells such as the Kupffer cells, dendritic cells, natural killer cells, etc., over 90% of the volume of the liver is made up of its parenchymal cells-the hepatocytes (57, 62, 63). Therefore how the hepatocytes respond to foreign antigens or infectious agents would greatly influence the overall immune portfolio of the liver (64). Hepatocytes limiting the extent of caspase-1 processing and pro-inflammatory cytokine responses while retaining the capacity to undergo pyroptosis may be a critical adaptation that maintains the immunotolerant nature of the liver without surrendering the ability to combat pathogens. Gaining a deeper mechanistic understanding of how various innate immune networks operate within the hepatocytes is necessary to harness the full immune potential of the liver and manage liver diseases such as those driven by inflammatory responses, like liver fibrosis, cirrhosis etc., and our work is a key step in this direction.

When caspase-1 processing was artificially limited to the p32 form by mutating the IDL sequence of procaspase-1, BMDMs were unable to efficiently mature IL-1β in response to LPS+ATP stimulation or *St* infection (11). Similarly, altering ASC sequence through site-directed mutagenesis restricted the efficient formation of inflammasome complexes in BMDMs, impeding the maturation of IL-1β (42). Nevertheless, these alterations did not prevent GSDMD activation or the ability of the host cells to undergo pyroptotic cell death. Notably, although less efficient in comparison to the fully processed caspase-1, procaspase-1 is capable of activating GSDMD and inducing cell death to a certain degree (42, 65). This meant that the proinflammatory role of caspase-1 can be regulated in cells by limiting the extent of procaspase-1 processing, and that various cell lineages may have ontogenically restricted or acquired the full range of functions offered by caspase-1 by altering the expression of inflammasome components such as the ASC adaptor. A variety of non-canonical caspase-1 cleavage products are observed in neurons (66, 67), epithelial cells (68), corneal stromal cells (69), or certain types of carcinomas (70–73). Various caspase-1 cleavage products are also visible in myeloid cells following infection or PAMP stimulation (74, 75). An intermediate product of procaspase-1 processing composed of unseparated CARD and p20 subunits, called p33 has been described in macrophages (27). In contrast to the long-lived p32 composed of p20 and p10 caspase-1 subunits, p33 in macrophages is composed of the CARD and p20 domains and is considerably short-lived (< 30 minutes) (2, 27). Only future studies will help determine the origins and biological relevance of such ‘non-conformist’ caspase-1 species observed in the different cell types.

The convention of undertaking biochemical and functional characterizations of inflammasome pathways using immune cells belonging to the myeloid lineage have also created the notion that pro-inflammatory cytokine responses and pyroptotic cell-death are inseparable consequences of caspase-1 activation (7, 76–78). Our observation that inflammasome-mediated caspase-1 activation in hepatocytes do not induce pro-inflammatory cytokines challenges this idea. We believe that the ability to process procaspase-1 into the p20 and p10 domains may have evolved with functional specialization in the immune cells of myeloid lineage. Increased availability of ASC in these cells would create a denser platform to bring procaspase-1 molecules in close proximity to each other and enable its complete proteolysis (42). Mutating ASC to impede their inter-molecular interaction generates larger, less condensed ASC specks in macrophages, resulting in significantly reduced caspase-1-mediated functions (42). We showed that diminishing ASC expression in BMDMs reduced the efficiency of caspase-1 cleavage, leading to the appearance of caspase-1p32, presumably at the cost of caspase-1p20. The increased basal expression of ASC may have enabled the myeloid cells to mature pro-inflammatory cytokines such as IL-1 and IL-18, thereby fulfilling their signaling and recruitment functions. A lower expression of ASC, on the other hand, would limit caspase-1 processing in parenchymal cells such as the hepatocytes, which are not professional immune cells. Intriguingly, neutrophils also express ASC at lower levels compared to macrophages and exhibit unconventional caspase-1 processing dynamics and function following PAMP stimulation or pathogen encounter (2, 27, 69, 79). Certain pathogens such as Sendai virus and paramyxoviruses are known to regulate inflammasome activation and caspase-1 processing dynamics in host cells by targeting ASC oligomerization (80). Human adenovirus 5 can inhibit ASC phosphorylation and oligomerization to evade inflammasome-mediated caspase-1 activation (81). Therapeutically impeding ASC oligomerization has been shown to result in broad-spectrum inhibition of inflammasome activation and is a proposed treatment option for inflammasome-dependent inflammatory disorders (82). Targeting ASC expression and function may therefore offer a potential path to alter the extent and consequences of the induction of inflammasome in the context of infectious or inflammatory diseases. We believe that the inherently reduced expression of ASC in hepatocytes may have played a significant role in the evolutionary choice of *Plasmodium* to undertake pre-erythrocytic development in these cells in mammals.

CSF1R^+^ CD11c^+^ APCs are instrumental in the generation of protective CD8 T cell responses following natural malaria infections and sporozoite-based anti-malarial vaccination (4). Conventional wisdom would dictate that enhancing the recruitment of such CSF1R^+^ APCs to the liver would therefore result in the generation of stronger immune responses and protection from future *Plasmodium* challenges. However, RAS vaccination in the context of ASC over-expression in hepatocytes resulted in diminished CD8 T cell responses and suboptimal protective immunity. Prior research has shown that prolonged survival of the hepatocytes harboring live-attenuated vaccine strains of *Plasmodium* is critical for the generation of robust immune responses against malaria (83). Adequate development and persistence of the attenuated parasites in hepatocytes is believed to made a broader range of protective epitopes available to the antigen-presentation machinery in the liver (13, 83). Over-expression of ASC in hepatocytes likely results in rapid clearance of the RAS-infected hepatocytes. Of note, intact type I IFN signaling in hepatocytes which facilitates such rapid clearance of *Plasmodium* from the liver, albeit via other potential mechanisms also limits the immune responses and protection from malaria following sporozoite-based vaccination (2, 3, 56). Therefore, therapeutically limiting ASC oligomerization and function further in the hepatocytes, such as with the drug MM01, may be a viable strategy to enhance the efficacy of live-attenuated sporozoites based vaccines against malaria (84). Our work, in addition to filling a fundamental knowledge gap in our understanding of the dynamics of the regulation of procaspase-1 processing in cells, offers a new path to modify the extent and consequences of inflammasome processing in cells.

## Acknowledgements

We want to thank the UGA CTEGD Flow Cytometry Core and the UGA CTEGD Sporocore staff for their contributions, as well as the other members of the Kurup lab for their support. We also acknowledge Dr. Kojo Mensa-Wilmott for giving access to fluorescence microscopy, and Dr. Ronald Etheridge for valuable feedback on the manuscript.

## Author contributions

CMdS, RPB, and SPK designed the experiments. CMdS and RPB performed the experiments. CMdS and SPK wrote the manuscript.

