## Supplemental figures for "Inherently reduced expression of ASC restricts caspase-1 processing in hepatocytes and promotes *Plasmodium* infection"

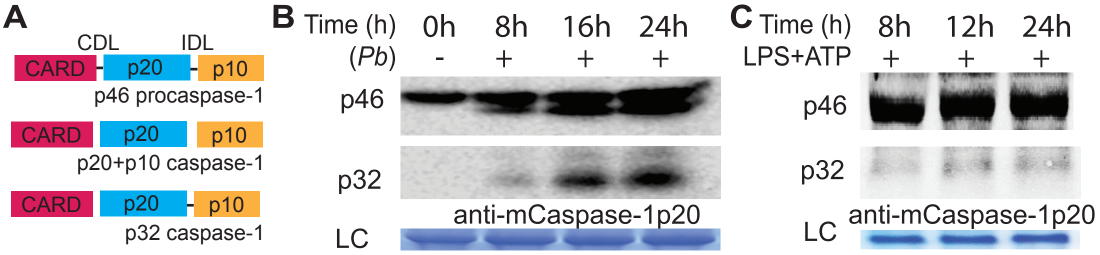
A
CDL
IDL
CARD
p20
p10
p46 procaspase-1
CARD
p20
p10
p20+p10 caspase-1
CARD
p20
p10
p32 caspase-1
B
Time (h)
0h
8h
16h
24h
(Pb)
-
+
+
+
p46
p32
anti-mCaspase-1p20
LC
C
Time (h)
8h
12h
24h
LPS+ATP
+
+
+
p46
p32
anti-mCaspase-1p20
LC

**FIGURE S1: Incomplete processing of caspase-1 in hepatocytes.**

(**A**) In the myeloid cells, autoproteolytic processing of the 46kDa procaspase-1 (p46) yields distinct CARD, p20 and p10 domains. The p20 and p10 domains subsequently associate with each other and generate a catalytically active hetero-tetramer, which is widely considered to be the stable, active form of caspase-1 (7). In the hepatocytes however, proteolysis of procaspase-1 yields a terminally processed p32 caspase-1 species composed of unseparated p20 and p10 domains (2). CDL: CARD domain linker, IDL: Interdomain linker.

(**B-C**) Immunoblot analysis for caspase-1 processing in primary mouse hepatocytes infected with *Pb* sporozoites (**B**) or treated with LPS+ATP (**C**) for the indicated time-frames. LC: loading control. Immunoblots in **B-C** representative of $\geq$ 3 replicate experiments including previously published ones (2).

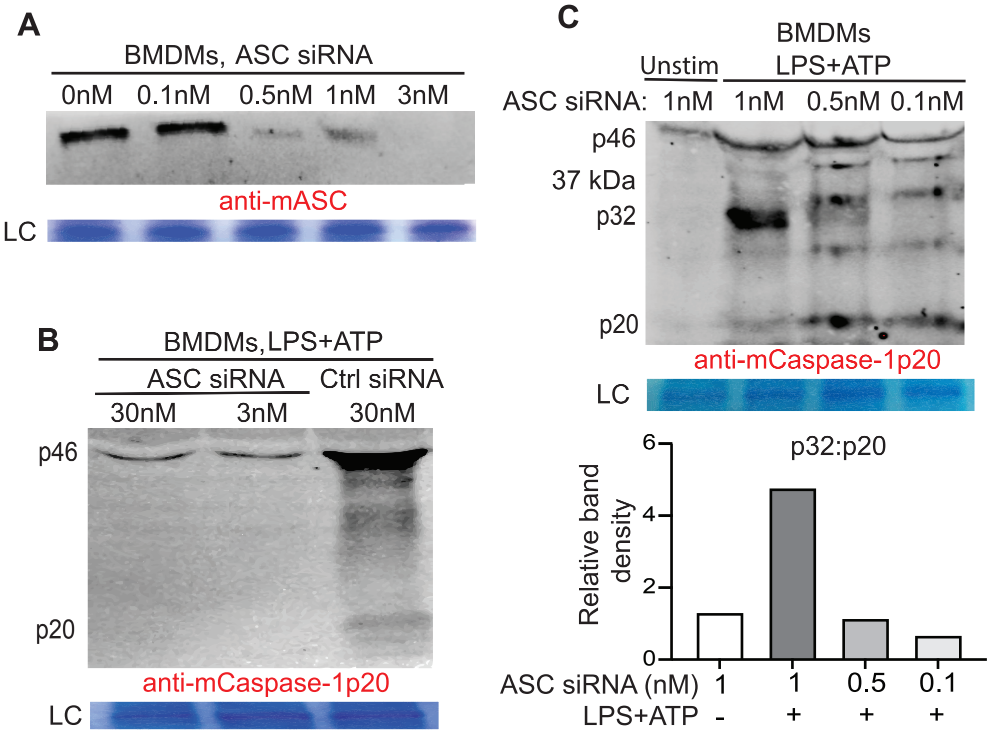
A
BMDMs, ASC siRNA
0nM 0.1nM 0.5nM 1nM 3nM
anti-mASC
LC
B
BMDMs,LPS+ATP
ASC siRNA
Ctrl siRNA
30nM 3nM 30nM
p46
p20
anti-mCaspase-1p20
LC
C
BMDMs
Unstim
LPS+ATP
ASC siRNA: 1nM 1nM 0.5nM 0.1nM
p46
37 kDa
p32
p20
anti-mCaspase-1p20
LC
p32:p20
Relative band density
0
2
4
6
ASC siRNA (nM) 1 1 0.5 0.1
LPS+ATP - + + +

**FIGURE S2. Reducing ASC expression engenders incomplete caspase-1 processing in BMDMs.**

(**A**) Immunoblot analysis depicting the relative expression of ASC in BMDMs isolated from B6 mice and transfected with the indicated dosages of ASC siRNA *in vitro*, at 24h post-transfection. Representative blot shown from 3 replicate experiments.

(**B**) Immunoblot analysis for caspase-1 processing in BMDMs transfected with the indicated doses of ASC siRNA or control scrambled (ctrl) siRNA, followed by stimulation with LPS (3.5h)+ATP (0.5h) from 24h post-transfection. Representative blot shown from 2 replicate experiments.

(**C**) Immunoblot analysis for caspase-1 processing in BMDMs transfected with diminishing dosages of ASC siRNA and stimulated with LPS+ATP as above at 24h post transfection. LC: loading controls. The bar graph below summarizes the ratio of the p32 and p20 band densities in the presented blot. Unstimulated cells with no detectable p32 or p20 subunits was given an arbitrary value of 1. A representative blot shown from 3 separate replicate experiments.
